# Zbtb14 regulates monocyte and macrophage development through inhibiting *pu.1* expression in zebrafish

**DOI:** 10.1101/2022.07.03.498625

**Authors:** Yun Deng, Haihong Wang, Xiaohui Liu, Hao Yuan, Jin Xu, Hugues de Thé, Jun Zhou, Jun Zhu

## Abstract

Macrophages and their precursor cells, monocytes, are the first line of defense of the body against foreign pathogens and tissue damage. Although the origins of macrophages are diverse, some common transcription factors (such as PU.1) are required to ensure proper development of monocytes/macrophages. Here we report that the deficiency of *zbtb14*, a transcription repressor gene belonging to *ZBTB* family, leads to an aberrant expansion of monocyte/macrophage population in zebrafish. Mechanistically, Zbtb14 functions as a negative regulator of *pu*.*1*, and SUMOylation on a conserved lysine is essential for the repression activity of Zbtb14. Moreover, a serine to phenylalanine mutation found in an acute myeloid leukemia (AML) patient could target ZBTB14 protein to autophagic degradation. Hence, *ZBTB14* is a newly identified gene implicated in both normal and malignant myelopoiesis.

## Introduction

Hematopoiesis is the process by which hematopoietic stem cells (HSCs) proliferate and differentiate into all blood lineages. It is driven by a variety of transcription factors, which function in a stage and lineage specific manner (***Bodine,2017***). A major goal to explore these transcription factors and their regulatory networks is to gain an intensive insight into normal hematopoiesis and its malignant counterpart, leukemia.

Macrophages are key players in many biological processes such as immune response to foreign pathogens and tissue homeostasis. The developmental origin of macrophages is diverse (***Kurotaki, et al.,2017***). Most tissue-resident macrophages arise from the blood islands of the yolk sac (***Kurotaki, et al.,2017***). Yet, monocytes in circulation are derived from HSCs, which give rise to monocytes in a step-wise manner via common myeloid progenitors (CMPs), granulocyte–monocyte progenitors (GMPs), monocyte–dendritic cell progenitors (MDPs), and common monocyte progenitors (cMoPs) (***Hettinger, et al.,2013***). During infection, circulating monocytes migrate into tissues and generate inflammatory macrophages (***Menezes, et al.,2016***). Whatever the origin, some common transcription regulators (such as PU.1) (***Glass and Natoli,2016***; ***Kueh, et al.,2013***) and signaling pathways (such as macrophage colony-stimulating factor receptor, M-CSFR) are required in monocyte and macrophage development (***Dai, et al.,2002***).

A total of 49 zinc finger and BTB domain (ZBTB) containing transcription factors have been identified in human genome. The C-terminal zinc finger motifs of ZBTB proteins enable the binding with DNA, whereas the N-terminal BTB motif mediates the homo/hetero-dimerization/multimerization between different ZBTB family members and recruits corepressors such as NCoR and SMRT (***Maeda,2016***). Thus, ZBTBs mostly function as transcription repressors.

Several members of ZBTB family transcription factors including ZBTB16 (also known as promyelocytic leukemia zinc finger, PLZF) (***Sobas, et al.,2020***), ZBTB27 (also known as B cell leukemia/lymphoma 6, BCL6) (***Nakamura,2000***), ZBTB7 (also known as leukemia/lymphoma-related factor, LRF) (***Constantinou, et al.,2019***), and ZBTB15 (also known as T-helper-inducing POZ/Krueppel-like factor, ThPOK) (***Taniuchi,2016***) play various roles in both normal and malignant hematopoiesis in humans. In addition, at least 12 *Zbtb* genes are involved in hematopoietic development in mice (***Maeda,2016***).

ZBTB14, a ZBTB family member whose function has been poorly characterized, is expressed in a variety of blood cell types (The Human Protein Atlas). Recently, a missense mutation of *ZBTB14* gene (*ZBTB14*^*S8F*^) was detected by whole-exome sequencing in a newly diagnosed acute myeloid leukemia (AML) patient (***Tyner, et al.,2018***). Note that mutation of *ZBTB14* has not previously been identified in AML. [Nevertheless, the potential role of *ZBTB14* in hematopoiesis and leukemogenesis was obscure](https://www.ncbi.nlm.nih.gov/pubmed/25583766).

In the present work, we provide *in vivo* evidence showing that the deficiency of *zbtb14* leads to an expansion of monocyte/macrophage population in zebrafish. Mechanistic studies reveal that Zbtb14 functions as a negative regulator of *pu*.*1*, and SUMOylation on a conserved lysine is essential for the transcriptional repression of Zbtb14. In addition, human ZBTB14^S8F^ mutant protein is demonstrated as a loss-of-function transcription factor. Hence, our results for the first time, not only unravel the physiological function of Zbtb14 during monocyte/macrophage development, but also elucidate the defective role of its mutant in AML.

## Results

### Generation of a *zbtb14*-deficient zebrafish line

The zebrafish serves as an ideal model organism for hematopoietic development and disease studies (***Gore, et al.,2018***). Zebrafish Zbtb14 protein shares 70% homology to its human counterpart ZBTB14. The two transcription factors bear a nearly identical N-terminal BTB motif and five consecutive C-terminal zinc finger motifs (Fig 1A).

**Fig 1.**
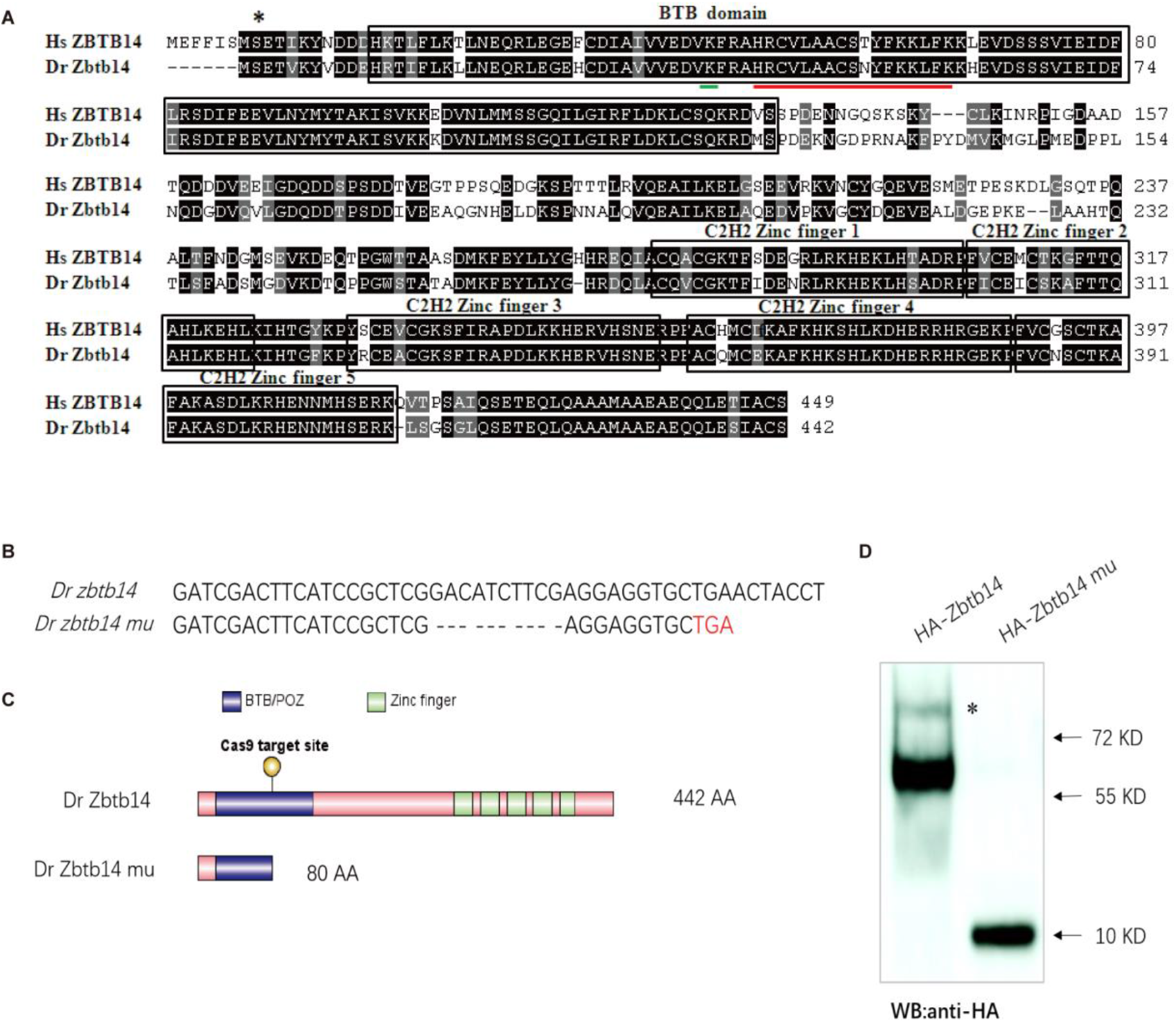
The establishment of a zebrafish *zbtb14* knockout line. (A) Sequence alignment of ZBTB14 and Zbtb14 proteins. Hs: homo sapiens, Dr: danio rerio. The conserved BTB domain and C2H2 zinc finger domains are boxed. *: The mutated serine identified in an AML patient. The putative SUMOylated lysine K40 and the nuclear localization signal (NLS) are underlined, respectively. (B) Schematic representation of Cas9 target site in the first exon of zebrafish *zbtb14*. The deleted nucleotides in the mutant gene are marked by hyphens. (C) Schematic representation of wild type (442 amino acids) and mutant Zbtb14 proteins (80 amino acids). (D) Western blot analysis of HA-tagged wild type and mutant Zbtb14 proteins expressed in HEK293 cells. * indicates the adduct band.

To explore the roles of *zbtb14* in hematopoiesis, especially in myelopoiesis, a mutant zebrafish line was established by using the CRISPR/Cas9 system targeting the BTB domain of Zbtb14. 10 nucleotides were deleted, which resulted in a truncated protein with only 80 amino acids by frameshifting (Fig 1B, C). The mutant *zbtb14* gene was expressed in HEK293T cells, and western blot analysis revealed a short protein as anticipated (Fig 1D). Note that the presence of a weak adduct, implying certain covalent modification of Zbtb14 protein might exist.

### Deficiency of *zbtb14* specifically affects monocyte and macrophage development

Like mammalian hematopoiesis, zebrafish hematopoiesis consists of primitive and definitive waves which occur sequentially in distinct anatomical sites (***Galloway and Zon,2003***; ***Zon,1995***). The primitive hematopoiesis gives rise to embryonic myeloid cells (neutrophils and macrophages) and erythrocytes from two intraembryonic locations, the rostral blood island (RBI) (the equivalent of mammalian yolk sac) and intermediate cell mass (ICM). The definitive HSCs which give rise to all blood lineages initiate within the ventral wall of the dorsal aorta (VDA) (a tissue analogous to the mammalian aorta/gonad/mesonephros, AGM), then translocate to the caudal hematopoietic tissue (CHT) (the equivalent of fetal liver) and colonize in kidney marrow (the equivalent of bone marrow) in adults (***Bertrand, et al.,2008***; ***Bertrand, et al.,2007***).

To unravel the role of *zbtb14* during embryonic hematopoiesis, whole-mount mRNA in situ hybridization (WISH) analyses were performed with multiple hematopoietic lineage-specific markers in *zbtb14*-deficient embryos and larvae. A significant increase of macrophage markers including *mfap4, csf1ra*, and *mpeg1*.*1* (***Meijer, et al.,2008***; ***Spilsbury, et al.,1995***; ***Zakrzewska, et al.,2010***) was observed from 19.5 hours post-fertilization (hpf) to 3 days post-fertilization (dpf) (Fig 2A-G’, H). This expanded macrophage population was further confirmed in *zbtb14*^*-/-*^//*Tg(mpeg1*.*1:eGFP)* larvae (Fig 2I-J’, K). It is worth noting that the z*btb14*-deficient macrophages could still migrate toward the wound as wild type ones did, implying their function remains intact (Fig 2L-M’). In addition, WISH with *apoeb* probe (a specific microglia marker) and neutral red (NR) vital dye staining revealed an expansion of microglia (the macrophages that reside in brain) in *zbtb14*-deficient larvae (Fig 2N-Q’, R).

**Fig 2.**
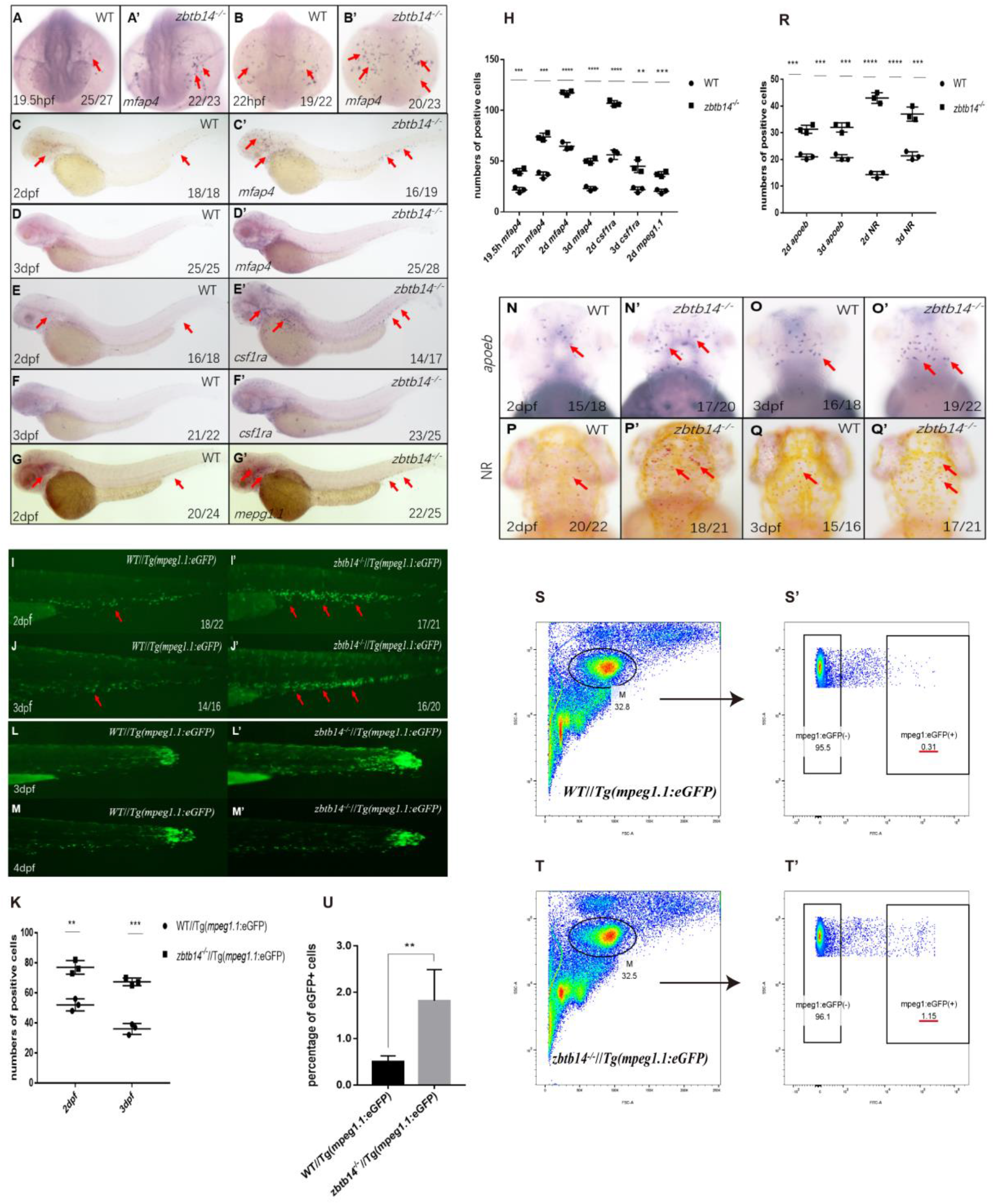
Deficiency of *zbtb14* specifically impairs monocyte and macrophage development in embryonic and adult zebrafish. (A-G’) WISH analyses of macrophage markers *mfap4* (A-D’), *csf1ra* (E-F’), *mpeg1*.*1* (G, G’) from 19.5 hpf to 3 dpf in wild type (WT) and *zbtb14*-deficient embryos and larvae. Red arrows indicate the main positions of positive cells for each marker. n/n, number of embryos/larvae showing representative phenotype/total number of embryos/larvae examined. (H) Statistical results for A-G’ (Student t test, N = 3, 14-28 embryos were used for each probe. Each dot represents the mean value of one experiment, which was obtained from the counts of all of the embryos/larvae in the same group. Error bars represent mean ± SEM. **P < 0.01, ***P <0.001, ****P < 0.0001). (I-J’) GFP positive cells were increased in *zbtb14*^*-/-*^ //*Tg(mpeg1*.*1:eGFP)* embryos at 2 and 3 dpf. (K) Statistical results for I-J’ (Student t test, N = 3, 14-22 larvae were used for each experiment. Each dot represents the mean value of one experiment. Error bars represent mean ± SEM. **P < 0.01, ***P <0.001). (L-M’) GFP positive cells in both *Tg(mpeg1*.*1:eGFP)* and *zbtb14*^*-/-*^//*Tg(mpeg1*.*1:eGFP)* larvae can migrate to the wound. (N-Q’) *apoeb* and neutral red positive cells were both increased in *zbtb14*-deficient larvae at 2 and 3 dpf. (R) Statistical results for N-Q’ (Student t test, N = 3, 15-22 larvae were used for each experiment. Each dot represents the mean value of one experiment. Error bars represent mean ± SEM. ***P < 0.001, ****P <0.0001). (S-T’) Representative scatterplot generated by FACS analysis of WKM samples collected from WT *Tg(mpeg1*.*1:eGFP)* (up panel) and *zbtb14-/-*//*Tg(mpeg1*.*1:eGFP)* (bottom panel) zebrafish lines in 4-month-old adults. M: myeloid gate. (U) Statistical results for S-T’ in wild type *Tg(mpeg1*.*1:eGFP)* and *zbtb14*^*-/-*^//*Tg(mpeg1*.*1:eGFP)* zebrafish. (Student t test, N = 4, each time 1 male and 1 female were used in the WT and mutant groups. Error bars represent mean ± SEM. **P < 0.01).

The primitive macrophages cannot sustain for a long period, which will eventually be replaced with HSC-derived macrophages as definitive hematopoiesis begins (***Xu, et al.,2012***). Since *zbtb14* mutant zebrafish were viable and fertile, the whole kidney marrow (WHM) samples were collected from wild type *Tg(mpeg1*.*1:eGFP)* and *zbtb14*^-/-^//*Tg(mpeg1*.*1:eGFP)* lines in adults. The myeloid cell population was analyzed, and many more macrophages were found in *zbtb14*^-/-^//*Tg(mpeg1*.*1:eGFP)* zebrafish than in controls (1.82% *mpeg1*.*1*^+^ vs 0.50% *mpeg1*.*1*^+^) (Fig 2S-T’, U), indicating that the development of macrophages originated from HSCs are also affected.

Nevertheless, the expressions of neutrophil and erythrocyte markers during primitive hematopoiesis stage, as well as those of hematopoietic stem and progenitor cells (HSPC), neutrophil, erythrocyte, and lymphocyte markers during definitive hematopoiesis stage were all normal (Fig S1). These observations suggest that only the development of monocyte/macrophage lineage is affected in the absence of *zbtb14*.

### Zbtb14 functions as a transcription repressor of *pu.1* in regulating monocyte and macrophage development

The monocyte/macrophage abnormalities in *zbtb14*-deficient larvae could be effectively rescued with either zebrafish *zbtb14* or human *ZBTB14* mRNA, confirming the specificity and conservation of ZBTB14 proteins (Fig 3A-D, F).

**Fig 3.**
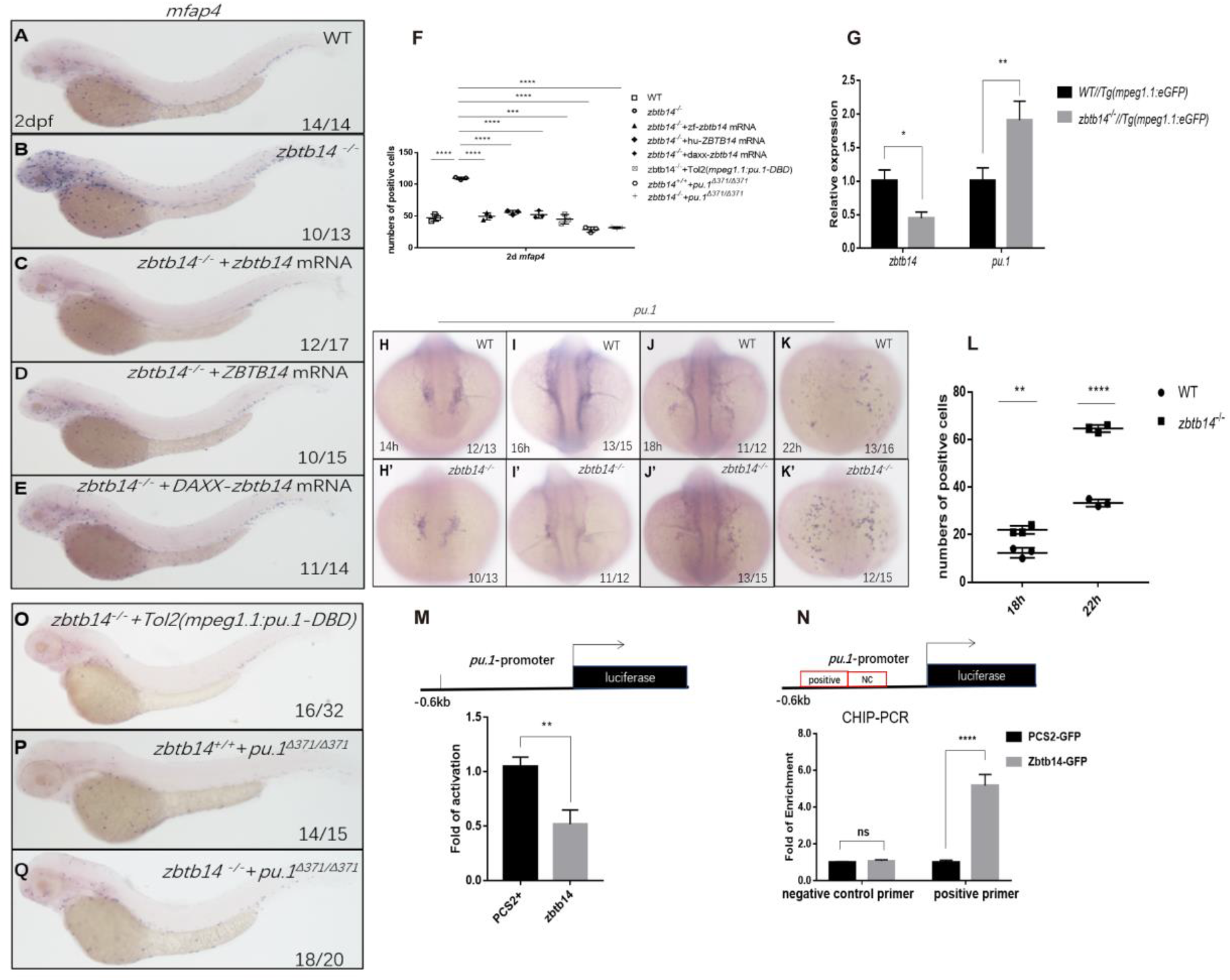
Zbtb14 regulates monocyte and macrophage development through inhibiting the expression of *pu*.*1*. (A-E) mRNA rescue assays in *zbtb14*^-/-^ larvae. *mfap4* probe was used in WISH to examine rescue effect with wild *zbtb14* (C), *ZBTB14* (D), *DAXX-ZBTB14* (E) mRNA injections. (F) Statistic result for A-E, and O-Q. The statistical significance was calculated by using one-way ANOVA. The statistical significance was calculated using 1-way ANOVA followed by Dunnett T3 correction. The asterisk indicates a statistical difference (N = 3, 10-32 larvae were used for each experiment. Each dot represents the mean value of one experiment. Error bars represent mean ± SEM. ****P < 0.0001). (G) Quantitative reverse transcriptase polymerase chain reaction analysis of *zbtb14* and *pu*.*1* in GFP positive cells enriched from *Tg(mpeg1*.*1:eGFP)* and *zbtb14*^*-/-*^//*Tg(mpx:eGFP)* larvae at 2 dpf. To determine the relative expression rate, data were normalized to the expression level of WT groups (which were set to 1.0) after normalized to the internal control of *β-actin* (Student t test, N = 3. Error bars represent mean ± SEM. *P < 0.1, **P < 0.01). (H-K’) Serial WISH analyses of *pu*.*1* in wild type and *zbtb14*^-/-^ embryos. (L) Statistical results for H-K’ (Student t test, N = 3, 10-16 embryos were used. Each dot represents the mean value of one experiment, which was obtained from the counts of all of the embryos in the same group. Error bars represent mean ± SEM. **P < 0.01, ****P <0.0001). (M) Luciferase reporter assay of Zbtb14 on the *pu*.*1* promoter. Bars showed the relative luciferase activity on the zebrafish *pu*.*1* promoter (−0.6 kb) (Student t test, N = 3. Error bars represent mean ± SEM. **P < 0.01). (N) CHIP-PCR analysis of *pu*.*1* promoter in zebrafish larvae expressing GFP or Zbtb14-GFP using an anti-GFP antibody. Positive: the location of the positive primers. NC: the location of the negative control primers. The statistical significance was calculated by using one-way ANOVA. The asterisk indicates a statistical difference (N =3. Error bars represent mean ± SEM. ns: not statistically significant, ****P < 0.0001). (O-Q) WISH assay of *mfap4* in *zbtb14*^*-/-*^ mutants injected with TOL2 *mpeg1*.*1*:*Pu*.*1 DBD, pu*.*1*^*Δ371/Δ371*^ mutants, and *zbtb14*^*-/-*^//*pu*.*1*^*Δ371/Δ371*^ double mutants.

ZBTB family proteins are frequently described as transcription repressors, nevertheless, ZBTB14 displays an activation or inhibition effect on different promoters (***Kaplan and Calame,1997***). To distinguish the activity of Zbtb14 on transcription, the repression domain of DAXX (a transcription corepressor) (***Zhou, et al.,2006***) was fused in frame with Zbtb14 (DAXX-Zbtb14), which forced the fusion protein to be a potent repressor. *In vivo* rescue assays demonstrated that *DAXX-zbtb14* mRNA had an obvious rescue effect as wildtype *zbtb14* mRNA (Fig 3E, F), implying Zbtb14 acted as a negative regulator in monocyte/macrophage development.

To elucidate the mechanism underlying the aberrant monocyte/macrophage development, we performed RNA sequencing (RNA-seq) analyses on *mpeg1*.*1*^*+*^ cells isolated from *Tg(mpeg1*.*1:eGFP)* and *zbtb14*^*-/-*^//*Tg(mpeg1*.*1:eGFP)* larvae at 2 dpf. The expression level of *pu*.*1*, the most critical transcription factor in monocyte/macrophage development, was obviously increased. Such upregulation was further confirmed by real-time quantitative PCR (RT-qPCR) analyses (Fig 3G). In addition, WISH analyses showed that the signals of *pu*.*1* were clearly intensified from 18 hpf to 22 hpf in *zbtb14*-deficient embryos compared with the wild type ones (Fig 3H-K’, L).

Since Zbtb14 was identified as a transcription repressor, we postulated that *pu*.*1* would be a major direct target of Zbtb14, whose derepression in the absence of Zbtb14 probably contributed to the expansion of monocyte/macrophage population. To test this hypothesis, a-0.6 kb zebrafish *pu*.*1* promoter in luciferase reporter was co-transfected with *zbtb14* expressing plasmid in HEK293T cells. As anticipated, luciferase analyses showed that Zbtb14 displayed a significant repression effect (Fig 3M).

Next, *in vivo* chromatin immunoprecipitation polymerase chain reaction (CHIP-PCR) was conducted in zebrafish larvae expressing GFP or GFP-Zbtb14 using an anti-GFP antibody. In this experiment, the *pu*.*1* promoter region could be specifically co-immunoprecipitated with GFP-Zbtb14 (Fig 3N).

To further demonstrate that *pu*.*1* was upregulated in macrophage lineage, a series of *in vivo* experiments was performed. A prominent rescue effect could be obtained with a dominant-negative *Pu*.*1* plasmid (the DBD domain of Pu.1 was under the control of *mpeg1*.*1* gene’s promoter, cloned in TOL2 backbone) injection in *zbtb14* mutants (Fig 3O, F). Moreover, we took advantage of a zebrafish *pu*.*1*^*Δ371*^ mutant line (a truncated Pu.1 whose transactivation activity was reduced) in which macrophages were reduced (***Yu, et al.,2017***) (Fig 3P, F), and no obvious alleviation could be found in *zbtb14*^*-/-*^//*pu*.*1*^*Δ371*^ double mutant zebrafish, indicating *zbtb14* was epistatic to *pu*.*1* (Fig 3Q, F).

In summary, these findings suggest Zbtb14 [regulates monocyte and macrophage development through inhibiting the expression of *pu*.*1*](https://pubmed.ncbi.nlm.nih.gov/33227819/).

### SUMOylation is essential for the transcription repression of Zbtb14

Post-translational modification plays important roles in regulating the functions of substrate proteins. Similar to ubiquitination, protein SUMOylation is catalyzed by a sequential enzymatic cascade including E1 (SAE1/SAE2), E2 (UBC9), and E3 in which SUMO (Small Ubiquitin-like MOdifier) molecules are covalently attached to lysine residues within substrate proteins (***Eifler and Vertegaal,2015***). The SUMOylation is generally associated with transcriptional repression through the recruitment of corepressors such as NCoR and SMRT (***Garcia-Dominguez and Reyes,2009***; ***Valin and Gill,2007***). We have mentioned above the presence of a weak adduct which was ∼ 10 kD (the size of one SUMO molecule) larger than the unmodified Zbtb14 protein (Fig 1D). Combing the fact that Zbtb14 was a repressor, we reasoned Zbtb14 would be a SUMOylated substrate.

To validate this hypothesis, *zbtb14*-expressing plasmid was transfected with or without *UBC9* and *SUMO1* in HEK293T cells. Western blot analyses showed that the adduct band became much more intensive in the presence of UBC9 and SUMO1 (Fig 4A, lane 1 and 2). Furthermore, immuno-coprecipitation (Co-IP) assays showed that SUMO1 molecules could be co-precipitated with Zbtb14 (Fig 4B, lane 1). These results indicated Zbtb14 could be SUMOylated in cells.

**Fig 4.**
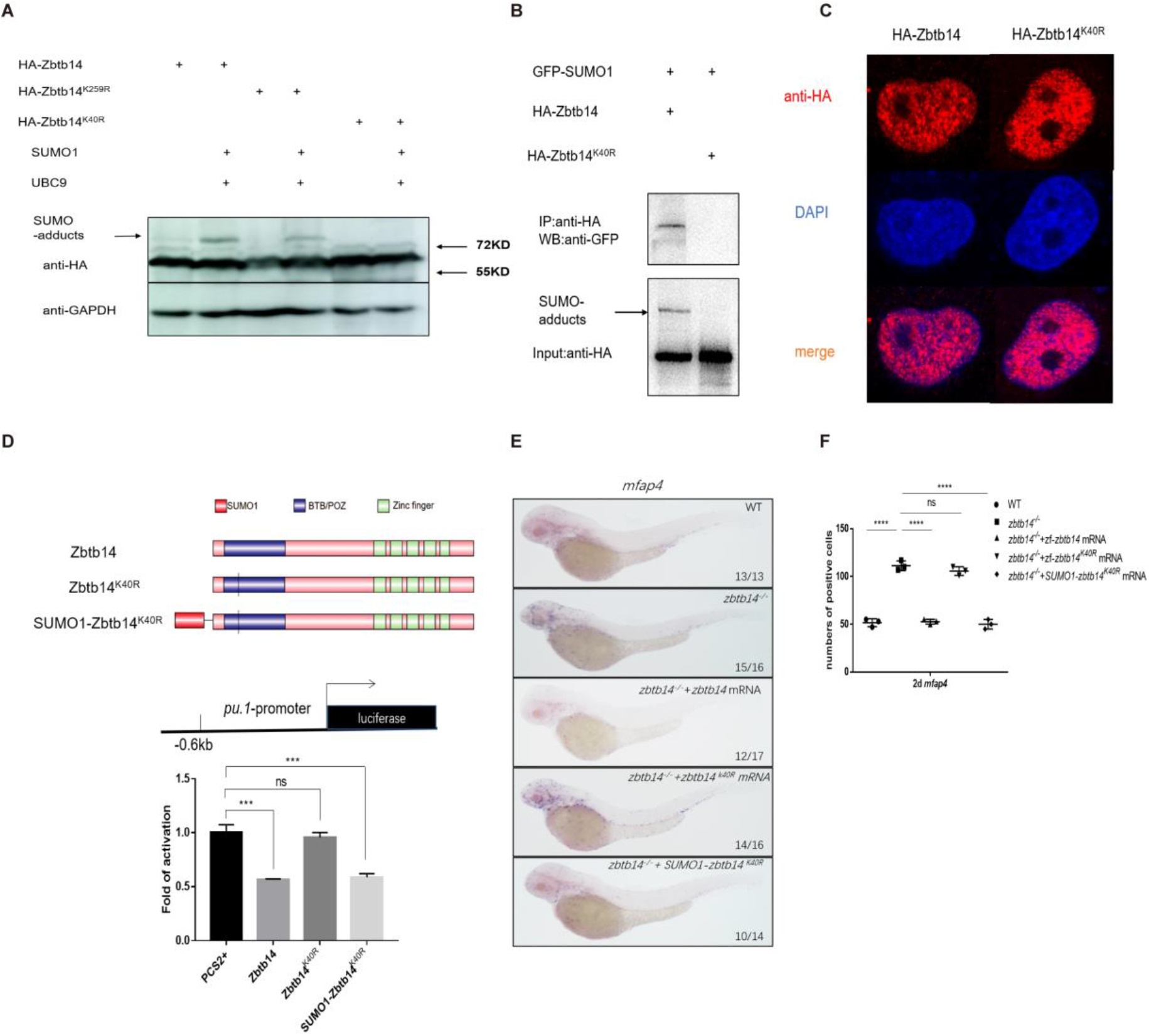
SUMOylation is indispensable for transcription repression of Zbtb14. (A) Western blot analysis (anti-HA) of HA-tagged wildtype (WT), Zbtb14^K40R^, and Zbtb14^K259R^ mutant proteins expressed in HEK293T cells in the absence or presence with the SUMO conjugating enzyme UBC9 and SUMO1. (B) HA-tagged WT or Zbtb14^K40R^ mutant protein was immunoprecipitated with an anti-HA antibody from HEK293T cells co-expressing GFP-SUMO1, and SUMOylated Zbtb14 protein was detected by western blot with an anti-GFP antibody. (C) Immunofluorescence analysis of wildtype (left panel) and Zbtb14^K40R^mutant protein (right panel). (D) The structure of variant forms of Zbtb14, including WT, Zbtb14^K40R^, and SUMO1-Zbtb14^K40R^ mutants (top panel). Repression of luciferase expression from the zebrafish *pu*.*1* promoter (−0.6kb) by Zbtb14 mutants (bottom panel). Bars showed the relative luciferase activity on the zebrafish *pu*.*1* promoter (−0.6 kb) (Student t test, N = 3. Error bars represent mean ± SEM. ns: not statistically significant, ***P < 0.001). (E) mRNA rescue assays in *zbtb14*^*-/-*^ mutant larvae. *mfap4* probe was used in WISH to examine rescue effects of injections of *zbtb14, zbtb14*^*K40R*^, and *SUMO1-zbtb14*^*K40R*^ mRNA. (F) Statistic result for E. The statistical significance was calculated by using one-way ANOVA. The asterisk indicates a statistical difference (N = 3, 10-17 embryos were used for each experiment. Each dot represents the mean value of one experiment. Error bars represent mean ± SEM. ns: not statistically significant, ****P < 0.0001).

SUMOylation is a process by which the SUMO monomer/polymer is covalently ligated to specific lysine residues of the target protein (***Chang and Yeh,2020***). Dozens of potential SUMOylation sites which spread throughout the protein were predicted by bioinformatics (SUMOsp2.0 prediction software, K259 has the highest score among 41 predicted SUMOylated sites). We carried out a series of mutations and finally found that the adduct band of Zbtb14^K40R^ mutant was abolished (Fig 4A, lane 3-6), suggesting K40 was the SUMOylated site.

Since K40 is located adjacent to the bipartite nuclear localization sequence (NLS) of Zbtb14 (***Sugiura, et al.,1997***) (Fig 1A), we questioned whether SUMOylation would affect its protein subcellular localization. HA-tagged Zbtb14 and Zbtb14^K40R^ were transfected in HEK293T cells, respectively. The results from immuno-fluorescence analyses revealed that the mutant protein was located in the nucleus as wildtype Zbtb14 (Fig 4C).

Next, luciferase reporter assays with the -0.6 kb zebrafish *pu*.*1* promoter were carried out. While Zbtb14 and SUMO1-Zbtb14^K40R^ (SUMO1 molecule was fused in frame with Zbtb14^K40R^ to mimic the SUMOylated Zbtb14) displayed a strong repression effect, Zbtb14^K40R^ completely lost the ability to repress transcription (Fig 4D).

Finally, *in vivo* rescue assays were conducted in *zbtb14*-deficient larvae with *zbtb14, zbtb14*^*K40R*^, and *SUMO1-zbtb14*^*K40R*^ mRNA, respectively. As expected, both *zbtb14* and *SUMO1-zbtb14*^*K40R*^ mRNAs had a remarkable rescue effect, whereas *zbtb14*^*K40R*^ mRNA did not (Fig 4E, F).

Taken together, these results suggest SUMOylation of Zbtb14 is pivotal for transcription repression.

### Human ZBTB14^S8F^ mutant is a loss-of-function transcription factor

A missense mutation, *ZBTB14*^*S8F*^, was detected in a *de novo* AML patient (***Tyner, et al.,2018***). To assess the role of the mutant protein, we first performed *in vivo* rescue assays in *zbtb14*-deficent zebrafish. While the wild type *ZBTB14* mRNA significantly rescued the expanded macrophage population, the mutant failed to display any rescue effect (Fig 5A, B), implying that the normal functions of ZBTB14 was lost.

**Fig 5.**
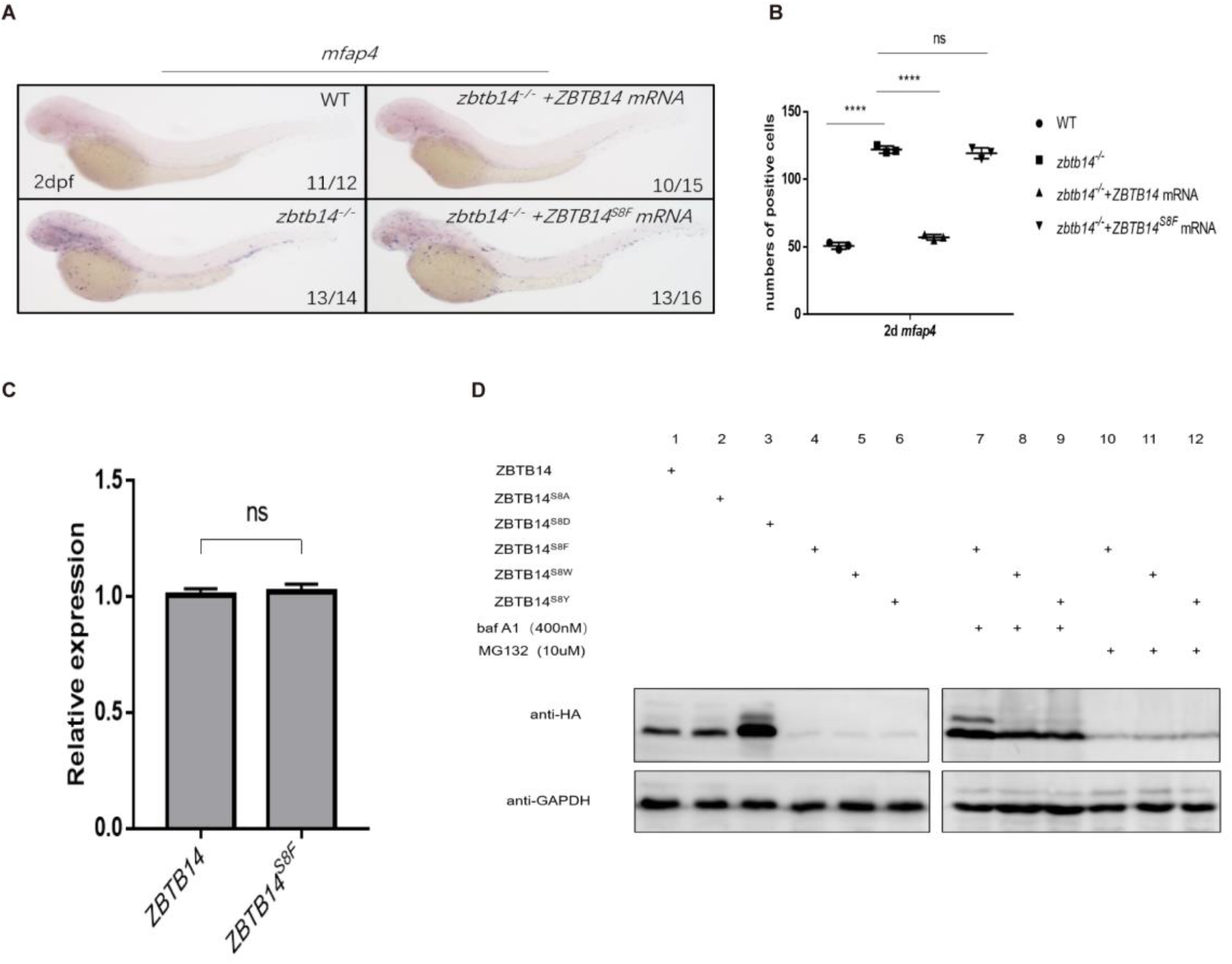
Human ZBTB14S8F mutant is a loss-of-function transcription factor. (A) mRNA rescue assays in zbtb14-/- mutant larvae. mfap4 probe was used in WISH to examine rescue effects of injections of ZBTB14 and ZBTB14S8F mRNA. (B) Statistic result for A. The statistical significance was calculated by using one-way ANOVA. The asterisk indicates a statistical difference (N = 3, 10-16 embryos were used for each experiment. Each dot represents the mean value of one experiment. Error bars represent mean ± SEM. ns: not statistically significant, ****P < 0.0001). (C) Quantitative reverse transcriptase polymerase chain reaction analysis of ZBTB14 and ZBTB14S8F transfected in HEK293T cells. To determine the relative expression rate, data were normalized to the expression level of WT groups (which were set to 1.0) after normalized to the internal control of β-actin (Student t test, N = 3. Error bars represent mean ± SEM. ns: not statistically significant). (D) Western blot analysis (anti-HA) of HA-tagged wildtype (WT), ZBTB14S8A, ZBTB14S8D, ZBTB14S8F, ZBTB14S8W, and ZBTB14S8Y mutant proteins expressed in HEK293T cells in the absence of presence with baf A1 or MG132. GAPDH served as internal control.

Next, we investigated the reason underlying the defects of the ZBTB^S8F^ mutant. Although its transcription level was comparable to that of wild type *ZBTB14* (Fig 5C), ZBTB14^S8F^ protein could hardly be detected by western blot analysis, implying that the stability of the mutant protein was impaired (Fig 5D, lane 4). We thus treated *ZBTB14*^*S8F*^ expressing HEK293T cells with MG132 (a proteasomal inhibitor), or bafilomycin (an autophagy inhibitor). The results indicated that bafilomycin, but not MG132, inhibited ZBTB14^S8F^ protein turnover (Fig 5D, lane 7 and 10), suggesting that the mutant protein might be subjected to autophagic degradation. Since serine is most frequently modified with phosphorylation, which could affect the protein stability, ZBTB14^S8A^ and ZBTB14^S8D^ mutants were constructed to mimic the dephosphorylated or constitutively phosphorylated status of ZBTB14. However, the turnover of both two mutant proteins still kept normal (Fig 5D, lane 2 and 3).

Autophagy is a process in which cellular material is degraded through the lysosomal pathway and recycled. Selective autophagy is mediated by specific cargo receptors which interact with the Atg8 (autophagy-related 8) family proteins and thereby link the cargo with the autophagy machinery (***Fracchiolla, et al.,2017***). MAP1LC3/LC3 (microtubule associated protein1 light chain 3) is one of the most important autophagy-related proteins which participates in autophagosome formation (***Fracchiolla, et al.,2017***). LC3 interacts with LIR (LC3-interacting region) motif (also termed as AIM, Atg8-interacting motif) of selective autophagy receptors that carry cargo for degradation. The LIR/AIM motif contains the consensus sequence [W/F/Y]xx[L/I/V] (***Popelka and Klionsky,2015***). The aromatic residue (W/F/Y) and hydrophobic residue (L/I/V) bind to two hydrophobic pockets formed by the ubiquitin-like fold of the Atg8 proteins (***Atkinson, et al.,2019***). Thus, the LIR/AIM ensures the selectivity of the interaction between Atg8 proteins and their binding partners. We noticed that the S to F mutation (<u>F</u>ET<u>I</u>) happened to form a φxxΨ motif, which was probably the reason why the ZBTB14^S8F^ mutant protein was targeted to undergo autophagic degradation. To further validate this point, ZBTB14^S8W^ and ZBTB14^S8Y^ mutants were also constructed and expressed in HEK293T cells. The results from western blots showed that the turnover of the two proteins was comparable to that of ZBTB14^S8F^ mutant protein (Fig 5D, lane 5, 6, 8, 9, 11, and 12).

Overall, the serine to phenylalanine mutation impairs the protein stability of ZBTB14, which probably contributes to AML pathogenesis.

## Discussion

In this study we characterize the roles of the previously enigmatic Zbtb14 in monocyte/macrophage development. Zbtb14 is absolutely required for proper monopoiesis during primitive hematopoiesis, which is reflected by the expansion of macrophage population in *zbtb14* mutant embryos/larvae compared to wild type ones from 19.5 hpf to 3 dpf. Such a requirement of Zbtb14 continues to definitive hematopoiesis since HSCs-derived macrophages remain hyperproliferative in *zbtb14*^*-/-*^ adults. Hence, *zbtb14* is indispensable for the maintenance of proper quantity of monocytes/macrophages. Functionally, the *zbtb14*-deficient *mpeg1*.*1*^+^ cells can still be recruited to the wound, and result in a normal healing, implying these macrophages are able to terminally differentiate into mature cells.

The mechanistic studies identify that *pu*.*1* is a direct target of Zbtb14. As a pivotal ETS family transcription factor, PU.1 is implicated in multiple stages of hematopoiesis such as generation of early myeloid progenitors, cell fate determination of granulocytic versus monocytic lineages of neutrophil-macrophage progenitors, maintenance of the accessibility of macrophage-specific genes during monocytic differentiation (***Kastner and Chan,2008***). We compared the expression level of *zbtb14* in *mpx*^+^ and *mpeg1*.*1*^+^ cells, and found *zbtb14* transcripts were much higher in the latter (Fig S2). This result together with the fact that the neutrophil population kept intact in *zbtb14* mutants, implying *zbtb14* is unnecessary for neutrophil lineage development.

Meanwhile, we also investigated the transcriptional network that involved *zbtb14*. Bioinformatic analyses show that PU.1 is a predicted upstream transcription factor of mammalian *ZBTB14*. The promoter region of zebrafish *zbtb14* was cloned into a reporter vector, and luciferase assay showed that PU.1 displayed a significant repression effect on it (Fig S3). Therefore, a Zbtb14-Pu.1 [negative feedback loop might regulate monocyte and macrophage development](https://pubmed.ncbi.nlm.nih.gov/33227819/).

Approximately 20 genes including *CEBPA, RUNX1, FLT3, DNMT3A*, and *NPM1* are most frequently mutated in AML patients (***Ley, et al.,2013***). Nevertheless, some rare mutations can be found in few samples (***Ley, et al.,2013***). A missense mutation, *ZBTB14*^*S8F*^, was detected in a *de novo* AML patient (***Tyner, et al.,2018***). In the current study, we took advantage of the *zbtb14*-deficient zebrafish line to demonstrate the functional conservation between the human *ZBTB14* and the zebrafish ortholog *zbtb14*. While wild type *ZBTB14* mRNA displayed a similar rescue effect as *zbtb14*, the *ZBTB14* mutant was shown to be a loss-of-function transcription regulator in monocyte/macrophage development. Furthermore, we demonstrated that the protein stability of ZBTB14^S8F^ was profoundly affected due to aberrant autophage.

*ZBTB14*, originally named as *ZF5*, was cloned as a transcriptional repressor gene on the murine *c-MYC* promoter (***Yanagidani, et al.,2000***). *C-MYC* amplification was identified in some AML patients (***Tang, et al.,2021***). Moreover, *c-MYC* is one of the most overexpressed genes in AML (***Handschuh, et al.,2018***). Either increased expression or aberrant activation of *c-MYC* plays important roles in leukemogenesis (***Salvatori, et al.,2011***). RNA-seq and RT-qPCR experiments showed that *c-myc* was upregulated in our *zbtb14*-deficient macrophages (Fig S4). In addition, it has been reported that *Znf161* (another alias of *Zbtb14*) knockout mice had a defect in genomic instability, which was associated with higher cancer risk (***Kim, et al.,2019***). These results suggest ZBTB14 would be a tumor suppressor, whose inactivity is tightly related with AML pathogenesis in humans.

## Materials and Methods

### Ethics Statement

The study was approved by the Ethics Committee of Rui Jin Hospital Affiliated to Shanghai Jiao Tong University School of Medicine. All animal work was approved by the Animal Care and Use Committee of Shanghai Jiao Tong University.

### Zebrafish maintenance and mutant generation

Zebrafish were raised, bred, and staged according to standard protocols (***Kimmel, et al.,1995***). The following strains were used: AB, *Tg(mpeg1*.*1:eGFP)* (***Ellett, et al.,2011***). For CRISPR9 mediated *zbtb14* knockout zebrafish generation, guide RNA (gRNA) targeting exon1 of *zbtb14* was designed using an online tool ZiFiT Targeter software (http://zifit.partners.org/ZiFiT), which was synthesized by cloning the annealed oligonucleotides into the gRNA transcription vector. Cas9 mRNA and gRNA were co-injected into one-cell stage zebrafish embryos. The injected F0 founder embryos were raised to adulthood and then outcrossed with wild type zebrafish. F1 embryos carrying potential indel mutations were raised to adulthood. Then PCR amplification and sequencing were performed on genomic DNA isolated from tail clips of F1 zebrafish to identify mutants.

### Whole-mount in situ hybridization (WISH)

Digoxigenin-labeled RNA probes were transcribed with T7, T3 or SP6 polymerase (Ambion, Life Technologies, Carlsbad, USA). WISH was performed as described previously (***Thisse and Thisse,2008***). The probes labeled by digoxigenin were detected using alkaline phosphatase coupled anti-digoxigenin Fab fragment antibody (Roche, Basel, Switzerland) with 5-bromo-4-chloro-3-indolyl-phosphate nitro blue tetrazolium staining (Vector Laboratories, Burlingame, CA, USA). 10 ∼ 30 embryos were used for each probe. The positive signals were counted under a microscope, and the mean value was obtained from the counts of all of the embryos in the same group.

### Neutral red staining

Zebrafish larvae were collected at indicated time and soaked in Neutral Red (2.5 mg/mL, Sigma-Aldrich, St Louis, MO, USA) overnight at 28.5 °C. Staining was then observed under a microscope.

### Cell collection and FACS analysis

Cell collection and FACS analysis were performed as described (***Traver, et al.,2003***). Wild type *Tg(mpeg1*.*1:eGFP)* and *zbtb14*^*-/-*^//*Tg(mpeg1*.*1:eGFP)* larvae were dissociated into single cells using 0.05% trypsin (Sigma-Aldrich, St. Louis, MO, USA) as previously described (***Yan, et al.,2015***). These dissociated cells were passed through a 40-μm mesh, centrifuged at 450g, and suspended in 5% FBS/PBS before addition of propidium iodide to a final concentration of 1 μg/ml for exclusion of dead cells. Wild type zebrafish (without GFP) were used as blank to determine the background values in GFP-controls. The GFP^+^cells of each group were collected from a total of ∼ 1000 larvae using a FACS Vantage flow cytometer (Beckton Dickenson) (∼ 300 larvae once, performed 3 times). For the whole kidney marrow (WKM) samples, FACS analysis was based on forward and side scatter characteristics, propidium iodide exclusion and GFP fluorescence. The GFP^+^ cells in the myeloid gate was enriched from WKM samples of wild type *Tg(mpeg1*.*1:eGFP)* and *zbtb14*^*-/-*^//*Tg(mpeg1*.*1:eGFP)* zebrafish (four-month-old, each time 1 male and 1 female were used in the WT and mutant groups).

### RNA sequencing and Quantitative RT-PCR

At 48 hpf, GFP positive cells were isolated from either wild type *Tg(mpeg1*.*1:eGFP)* or *zbtb14*^*-/-*^//*Tg(mpeg1*.*1:eGFP)* larvae by FACS. mRNA was extracted from sorted cells using RNeasy Micro (Qiagen, Manchester, UK) and mRNA libraries were constructed using NEBNext Ultra RNA Library Prep Kit for52 Illumina and sequenced under Illumina HiSeq X Ten with pair end 150bp (PE150).

The quantitative PCR was carried out with SYBR Green Real-time PCR Master Mix (TOYOBO, Osaka, Japan) with ABI 7900HT real-time PCR machine and analyzed with Prism software. *β-actin* was served as the internal control. The primers used are listed in Table 1. Each time a different batch of samples was used. The expression levels of each interested gene were normalized to internal control *β*-*actin* by real-time qPCR and compared with WT group which was set to 1.0. Real time qPCR was performed with gene specific primers and gene expression levels were analyzed by comparative CT method.

**Table 1.**
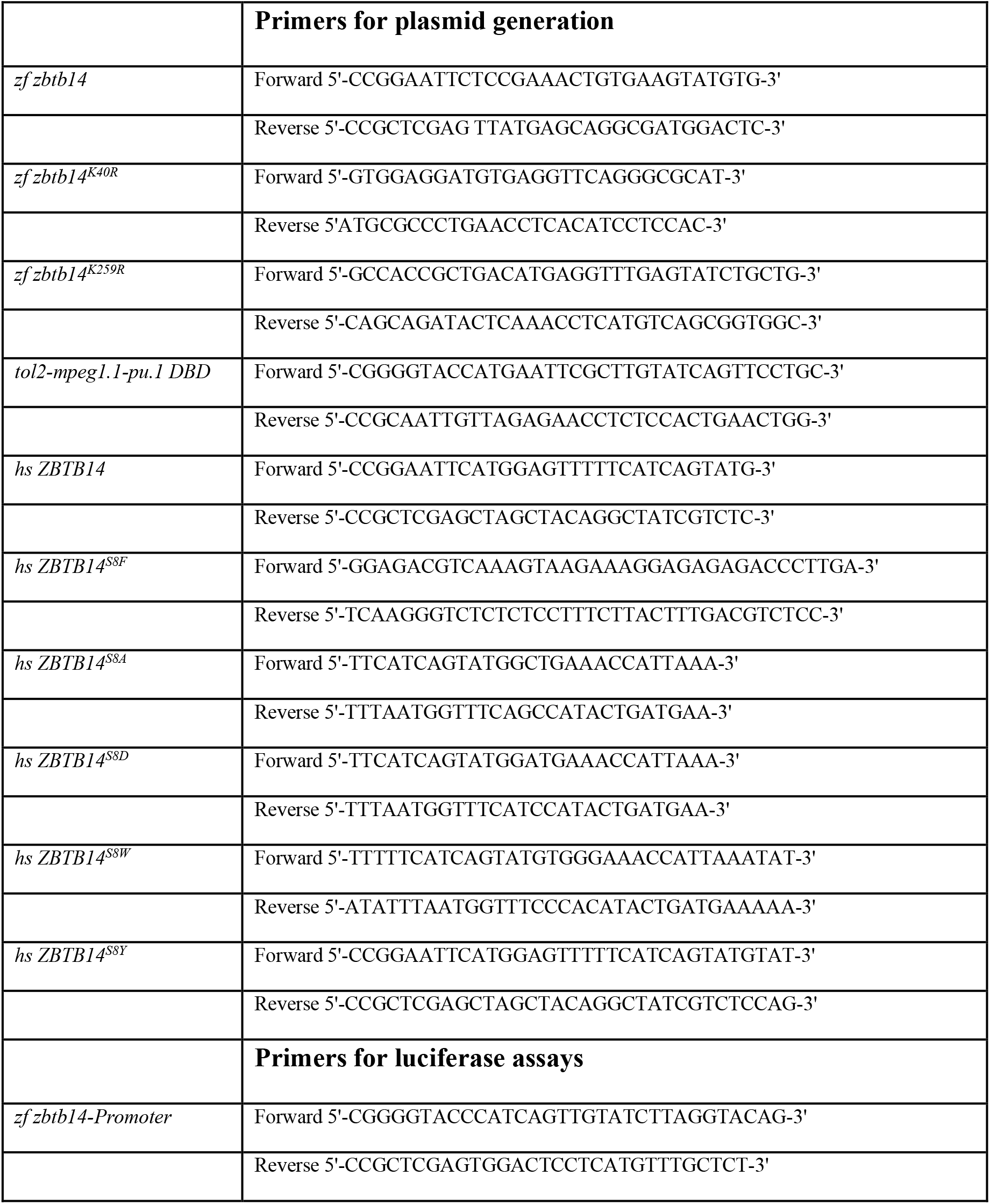
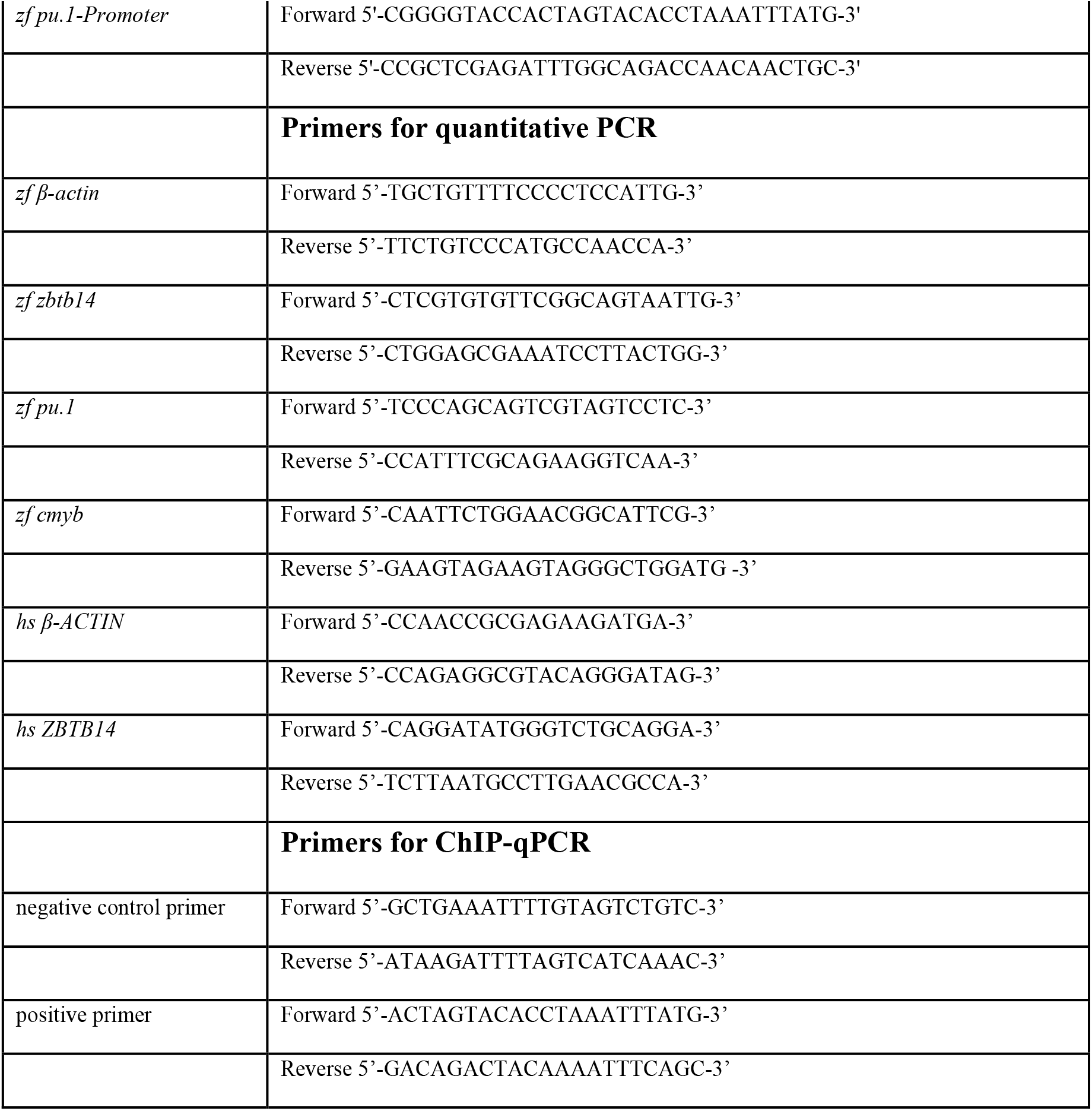
Primers for plasmid generation, luciferase assays, quantitative PCR, and CHIP-qPCR.

### Plasmid construction

Zebrafish *zbtb14* gene and its serial mutants were cloned into PCS2^+^ vector. For the luciferase reporter, the -0.6 kb promoter of zebrafish *pu*.*1* gene and the -1.1 kb promoter of zebrafish *zbtb14* gene were cloned into the PGL3 basic vector (Promega, Madison, WI, USA). Tol2-plasmid was constructed by insertion of *pu*.*1* DN under *mpeg1*.*1* promoter (2 kb). Transgene was transiently expressed by co-injecting 80 pg of Tol2-plasmid and 120 pg of Tol2 transpose mRNA at one-cell stage. Primers used were listed in Table 1.

### Morpholino and mRNA synthesis for microinjection

Zebrafish *zbtb14* (5’-ACTTCACAGTTTCGGACATACTGGA-3’),*pu*.*1* (5’-AATAACTGATACAAACTCACCGTTC-3’) targeting the transcriptional initiation ATG of *zbtb14, pu*.*1* was designed and purchased from Gene Tools. Full-length capped mRNA samples were all synthesized from linearized plasmids using the mMessage mMachine SP6 kit (Invitrogen, Thermo Fisher, Waltham, USA). Microinjection concentration of mRNA was between 50 ∼ 200 ng/μl and 2 nl of mRNA was injected at one-cell stage embryos. All injections were performed with a Harvard Apparatus micro-injector.

### Cell culture and luciferase reporter assay

HEK293T cells were maintained in DMEM (Gibco, Life technologies, Carlsbad, USA) with 10% Fetal Bovine Serum (Gibco, Life technologies, Carlsbad, USA). Plasmid transfection was carried out with Effectene Transfection Reagent (Qiagen, Manchester, UK) according to manufacturer’s instruction. For the luciferase reporter assay, cells were harvested 48 hours after transfection and analyzed using the Dual Luciferase Reporter Assay Kit (Promega, Maddison, WI, USA), according to the manufacturer’s protocols. Primers used were listed in Table 1.

### Chromatin immunoprecipitation PCR (ChIP-PCR)

For ChIP analysis, GFP and GFP-Zbtb14 expressing larvae were harvested at 48 hpf for brief fixation. Cross-linked chromatin was immunoprecipitated with anti-GFP antibody according to the procedure described (***Hart, et al.,2007***). The resultant immunoprecipitated samples were subjected to quantitative PCR using primer pairs (Table 1).

### Statistical analysis

Data were analyzed by SPSS software (version 20) using two tailed Student t test for comparisons between two groups and one-way analysis of variance (ANOVA) among multiple groups. Differences were considered significant at P<0.05. Data are expressed as mean ± standard error of the mean (SEM).

## Acknowledgments

The authors are grateful to Y Chen and J Jin (both from Shanghai Jiao Tong University School of Medicine, Shanghai, China) for technical support. We thank Dr. X Jiao (from the Department of Cell Biology and Neuroscience, Rutgers University, Piscataway, NJ 08854, USA) for his critical manuscript reading.

This work was supported by research funding from the National Natural Science Foundation of China (NO.32171097).

## Data availability

RNA sequencing dataset generated in this study was deposited with Dryad-https://doi.org/10.5061/dryad.9cnp5hqms.

