## Supplemental figures and figure legends for "Zbtb14 regulates monocyte and macrophage development through inhibiting *pu.1* expression in zebrafish"

### Figure S1

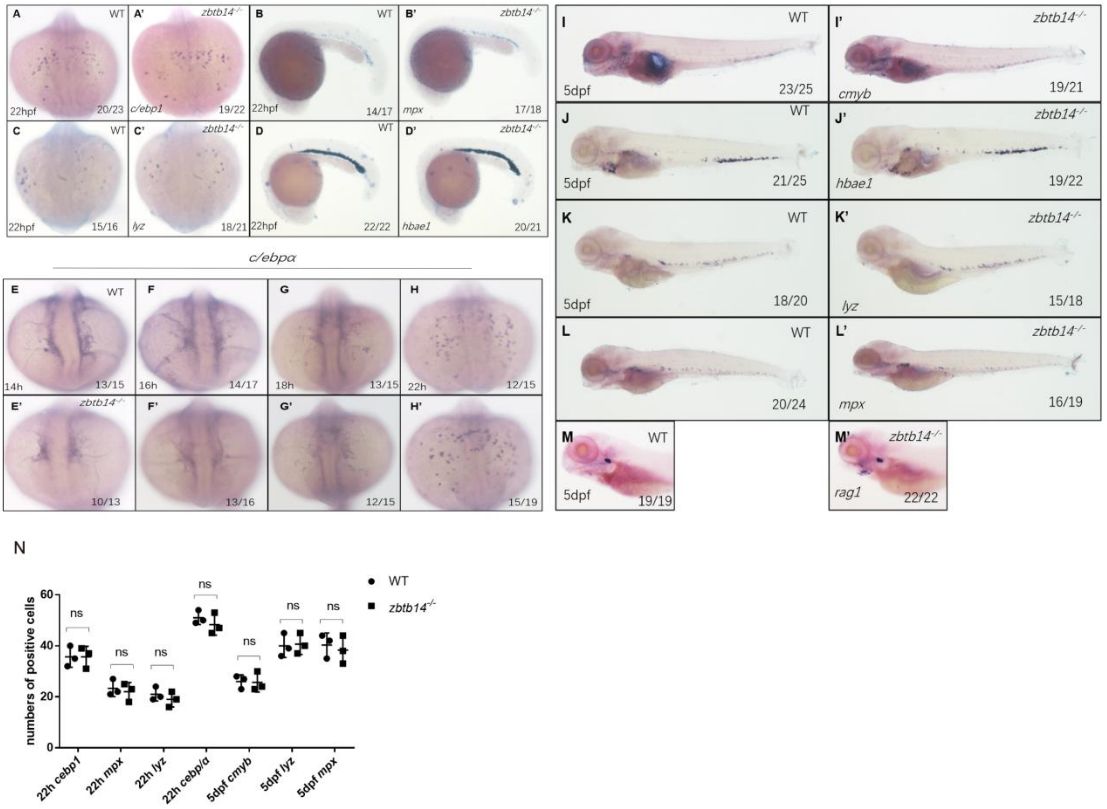


**Fig S1. Expression of lineage specific markers during primitive and definitive hematopoiesis stages in *zbtb14*-deficient embryos.** WISH analyses of neutrophil markers *c/ebp1*, *mpx*, *lyz*, and erythroid marker *hbae1* at 22 hpf (A-D'). WISH analyses of myeloid marker *c/ebpα* from 14 hpf to 22 hpf (E-H'). WISH analyses of HSPCs marker *c-myb*, erythroid marker *hbae1*, neutrophil markers *mpx*, *lyz*, and lymphoid marker *rag1* at 5 dpf (I-M'). (N) Statistical results for A-C', H-H', I-I', K-L' (Student t test, N = 3, 10-25 embryos or larvae were used for each probe. Each dot represents the mean value of one experiment, which was obtained from the counts of all of the embryos or larvae in the same group. ns: not statistically significant).

**Figure S2**

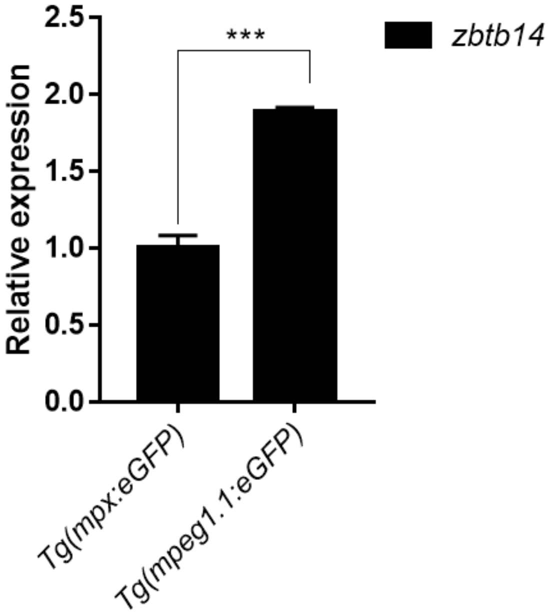


**Fig S2. Expression of *zbtb14* transcript in *mpx*$^{+}$ and *mpeg1.1*$^{+}$ cells.** Quantitative reverse transcriptase polymerase chain reaction analysis of *zbtb14* in GFP positive cells enriched from *Tg(mpx:eGFP)* and *Tg(mpeg1.1:eGFP)* larvae at 2 dpf. To determine the relative expression rate, data were normalized to the expression level of WT groups (which were set to 1.0) after normalized to the internal control of *β-actin* (Student t test, N = 3. Error bars represent mean ± SEM. ***P < 0.001).

**Fig S3**

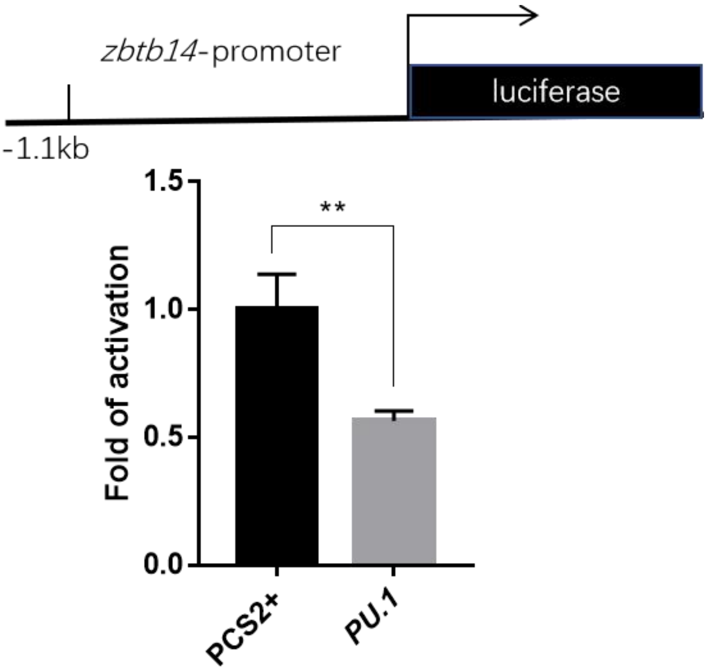


**Fig S3. Luciferase reporter assay of Pu.1 on the *zbtb14* promoter.** Bars showed the relative luciferase activity on the zebrafish *zbtb14* promoter (-1.1kb) (Student t test, N = 3. Error bars represent mean ± SEM. **P < 0.01).

**Fig S4**

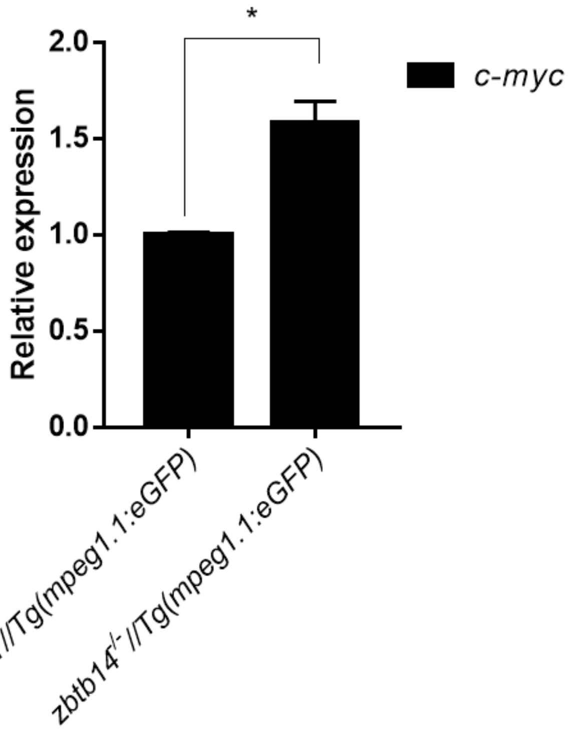


**Fig S4. Expression of *c-myc* transcript in *mpeg1.1*$^{+}$ cells.** Quantitative reverse transcriptase polymerase chain reaction analysis of *c-myc* in GFP positive cells enriched from *Tg(mpeg1.1:eGFP)* and *zbtb14*$^{-/-}$*//Tg(mpx:eGFP)* embryos at 2 dpf. To determine the relative expression rate, data were normalized to the expression level of WT groups (which were set to 1.0) after normalized to the internal control of *β-actin* (Student t test, N = 3. Error bars represent mean ± SEM. *P < 0.1).
